# Mitochondrial genome fragmentation is correlated with increased rates of molecular evolution

**DOI:** 10.1101/2023.06.22.546119

**Authors:** Tomáš Najer, Jorge Doña, Aleš Buček, Andrew D. Sweet, Oldřich Sychra, Kevin P. Johnson

**Affiliations:** Department of Veterinary Sciences, Faculty of Agrobiology, Food, and Natural Resources, Czech University of Life Sciences Prague, Prague 6, Czechia; Illinois Natural History Survey, Prairie Research Institute, University of Illinois Urbana-Champaign, Champaign, IL, USA; Departamento de Biología Animal, Universidad de Granada, Granada, Spain; Biology Centre, Czech Academy of Sciences, České Budějovice, Czechia; Department of Biological Sciences, Arkansas State University, Jonesboro, AR, USA; Department of Biology and Wildlife Diseases, Faculty of Veterinary Hygiene and Ecology, University of Veterinary Sciences Brno, Brno, Czechia

**Keywords:** mitochondrial genome, Psocodea, Phthiraptera, Amblycera, phylogenomics

## Abstract

Animal mitochondrial genomes (mitogenomes) typically exhibit a highly conserved gene content and organisation, with genes encoded on a single circular chromosome. However, many species of parasitic lice (Insecta: Phthiraptera) are notable exceptions, having mitogenomes fragmented into multiple circular chromosomes. To understand the process of mitogenome fragmentation, we conducted a large-scale genomic study of a major group of lice, Amblycera, with extensive taxon sampling. Analyses of the evolution of mitogenome structure across a phylogenomic tree of 90 samples from 53 genera, revealed evidence for multiple independent origins of mitogenome fragmentation, some inferred to have occurred less than five million years ago. We leveraged these many independent origins of fragmentation to compare the rates of DNA substitution and gene rearrangement, specifically contrasting branches with fragmented and non-fragmented mitogenomes. We found that lineages with fragmented mitochondrial genomes had significantly higher rates of mitochondrial sequence evolution. In addition, lineages with fragmented mitochondrial genomes were more likely to have mitogenome gene rearrangements than those with single-chromosome mitochondrial genomes. By combining phylogenomics and mitochondrial genomics we provide a detailed portrait of mitogenome evolution across this group of insects with a remarkably unstable mitogenome structure, identifying processes of molecular evolution that are correlated with mitogenome fragmentation.

**Author Summary:** Mitochondria are organelles that play a key role in providing energy to cells essential for life. The structure of the mitochondrial genome is conserved across most animal groups, being a single circular chromosome containing 37 genes. Deviations from this structure are typically detrimental and associated with some human diseases. However, in very few animal groups, the mitochondrial genome is fragmented into multiple circular chromsomes. In one group of insects, parasitic lice, fragmentation varies among species, with some having a complete circular genome and others having their mitochondrial genome fragmented in two or more smaller chromosomes. Here, we use whole genome sequencing reads to analyze an unprecedented number of species from a diverse group of lice (Amblycera) that exhibits both single-chromosome and fragmented mitochondrial genomes to understand how this fragmentation evolved. We found that fragmentation evolved many times independently in this group and this fragmentation is correlated faster rates mitochondrial molecular evolution and with an increased frequency of gene rearrangement. We also provide evidence that the rate of mitochondrial genome fragmentation changes over time. Altogether, our combination of broad sampling and phylogenomic and comparative analyses provide new insights into the mechanisms and dynamics of mitochondrial genome fragmentation.

## Introduction

Mitochondria play a vital role in the metabolism of eukaryotic organisms, supplying cells with most of the energy necessary to function. Likewise, the gene content, structure, and organisation of the mitochondrial genome (mitogenome) remain remarkably stable, especially across animals. In the animal kingdom, mitochondrial genetic information is usually organised on a single circular chromosome which is between 12,000–18,000 base pairs long and contains 37 genes [1]. Any defect in this structure typically results in cell death [2], and is linked to degenerative diseases, ageing, and cancer [3].

However, in some animals, mitogenome structure deviates from this conserved state without having a notable effect on mitochondrial functions. Specifically, the mitogenomes of some nematodes (*Globodera*; [4]), cnidarians [5], thrips [6], and lice [7] diverge from this typical structure. In all these groups, the original single mitochondrial chromosome is fragmented into smaller chromosomes of variable size and number, either linear or circular. Similar mitochondrial mutations have detrimental effects in humans [3, 8]. Therefore, understanding mitogenome fragmentation and reorganisation may provide significant insights for research on cell ageing or severe hereditary diseases.

Lice (Insecta: Psocodea) exhibit the most extensive variability in mitogenome structure among all animals. In free-living lice (i.e., bark lice), the mitogenomes are usually single-chromosome, except for book lice, which have two circular mitochondrial chromosomes [9]. In contrast, parasitic lice (Phthiraptera) show a broad range of mitogenome arrangements. Several lineages of parasitic lice maintain mitogenomes on single circular chromosomes [10], while in others, the mitogenomes are highly fragmented and consist of up to 20 small circular chromosomes [11, 12, 13]. Recent evidence of heteroplasmy of mitochondrial genome structure (presence of multiple structural variants within a single individual) in some lice [7] suggests that the initial fragmentation in lice is not a final state but an ongoing process towards increasing fragmentation. Hence, lice offer an exceptional model to document the process of mitogenome fragmentation over time.

One important question is whether the rate of mitogenome fragmentation might change over time. To date, studies of mitogenome fragmentation in lice have primarily focused on mammalian lice (Trichodectera; [14]; Anoplura; [13]; Rhynchophthirina; [15]) and avian feather lice (Ischnocera; [7, 11]). In the case of mammalian lice, the groups Anoplura (sucking lice), Rhynchophthirina (chewing lice), and Trichodectera (chewing lice) represent extreme instances of mitochondrial fragmentation. All members of these groups appear to have highly fragmented mitogenomes, with no cases of a single chromosome discovered in these groups to date. Thus, in Anoplura, Rhynchophthirina, and Trichodectera, mitochondrial fragmentation is highly phylogenetically conserved. However, in avian feather lice (Ischnocera), the opposite appears to be the case [7]. Within this group, mitogenome structure varies dramatically over the tree, with some species possessing a single mitochondrial chromosome while related genera are highly fragmented. This pattern makes it challenging to understand the process of stepwise fragmentation and whether there is phylogenetic conservation of mitogenome structure in Ischnocera. Given that mammalian lice and avian feather lice began diversifying around the same time [16], these differences in mitogenome structure suggest the rate of fragmentation might change over time across all parasitic lice. Mitochondrial fragmentation in another major group of lice (Amblycera) is less well studied. One study of nine species in this group revealed five cases of single-chromosome mitogenomes and four of fragmented mitogenomes, which seem to have occurred multiple times [10]. A more recent study provided further evidence of multiple fragmentation events in two families of Amblycera (Laemobothriidae and Menoponidae; [17]). Given the potential for multiple independent origins of mitogenome fragmentation, Amblycera provides an excellent opportunity to study the evolutionary dynamics of mitochondrial fragmentation with greatly expanded sampling and comparative analyses.

There are two principal, although not necessarily mutually exclusive, hypotheses for the origin and mechanism of mitochondrial fragmentation. The first is that mitochondrial fragmentation is related to errors in recombination within the mitochondrial genome [13, 18], even though it is not necessarily clear how this recombination might mechanistically occur. A second hypothesis for the origin of fragmentation suggests that initial heteroplasmy, in which a second partial copy of the mitochondrial genome is produced through replication errors, triggers the process of fragmentation [7]. Heteroplasmy of chromosome structure has been documented in lice with fragmented [7] and non-fragmented [12] mitogenomes. Heteroplasmy results in two (or more) copies of several genes and knockout mutations of one of the alternate copies could occur without being deleterious. If this happens, both chromosomes that contain functioning copies of different genes would then be preserved by selection and then this process could iterate leading to increasing fragmentation.

Both of these hypotheses have implications for the process of mitochondrial gene rearrangements, which are also widely observed in lice [19]. A single mitochondrion often has multiple copies of the mitogenome [20], which could facilitate recombination among the copies. Thus, recombination in the mitochondrial genome could cause gene order rearrangements. In comparisons across several metazoan phyla, Feng et al. [18] observed that lineages that had cases of mitochondrial fragmentation also exhibited higher rates of gene rearrangement; however, the level of rearrangement within clades was not related to whether mitogenomes were fragmented or not. Under the heteroplasmy hypothesis, gene order could also change because it is expected that genes with any knockout mutations would eventually be deleted in one of the heteroplasmic copies, which would create novel gene boundaries. Thus, both hypotheses predict that lineages with fragmented mitochondrial genomes should have increased rates of gene rearrangement.

Another feature that might be predicted under both hypotheses for mitochondrial fragmentation is an increased substitution rate and change in base composition of mitochondrial DNA [21]. Feng et al. [18] observed that mitochondrial substitution rates appeared to be faster in lineages that contained representatives with mitochondrial fragmentation. Across all animals, mitochondrial genomes typically exhibit a strong AT bias, often over 70% [7]. This phenomenon is most readily explained by biases during replication, where the separated DNA strands are exposed to deamination mutations [22]. Mitochondria lack the extensive repair mechanisms of nuclear genomes [23], making mitogenomes more susceptible to mutations. However, animals with fragmented mitogenomes consistently show less AT biased mitogenomes [7, 10]. One explanation for this pattern is that when a mitogenome becomes fragmented, the chromosomes are shorter and take less time to replicate. Consequently, less time is spent in a single-stranded state, perhaps making the genome less vulnerable to deamination mutations. Thus, in fragmented mitogenomes, deamination mutations would be less common and the entire mitogenome become less AT biased [7]. However, if there is some equilibrium AT content, which seems likely given that AT content tends to be conserved within a taxon [24, 25], then changes in the equilibrium AT content (via changes in frequency of deamination mutations) could result in higher apparent substitution rates in lineages with fragmented mitogenomes. Given that this change in AT content would be expected in lineages with fragmented mitogenomes, both hypotheses for the origins of fragmentation would predict this increased apparent substitution rate. In addition, it has been suggested [18] that lineages with increased rates of mutation might be more prone to recombination errors among mitochondrial genome copies, which could also lead to a correlation between substitution rate and mitochondrial fragmentation.

One pattern that might distinguish between the recombination model and heteroplasmy model is whether there are changes in the rate of fragmentation. In lineages where there is a complete mitochondrial genome, but also heteroplasmy of a partial genome, this model would predict a “chain reaction” in which fragmentation becomes much more likely. In contrast, in lineages that have no heteroplasmy, it is expected these lineages would be resistant to fragmentation. Under the recombination model, there is not necessarily an expectation that some lineages are more prone to recombination than others, and thus the rate of fragmentation might be predicted to be uniform across the tree.

The main goal of this study was to uncover the pattern of mitogenome fragmentation in a major group of lice, Amblycera, with unprecedented taxonomic and temporal resolution. Using genomic sequencing reads, we assembled the mitochondrial genomes of 90 samples of Amblycera, representing 53 genera and 89 species, and reconstructed a dated phylogenomic tree based on over 2,000 nuclear single-copy ortholog genes. With these data, we traced the phylogenetic pattern of the fragmentation across Amblycera, inferring the number of transitions from non-fragmented to fragmented mitogenomes and shedding light on the dynamics of this process. Furthermore, we used the evolutionary changes in fragmentation to estimate whether the rate of fragmentation changes across the tree. We also tested two key predictions of both models of mitogenome fragmentation by examining the correlation both gene rearrangement and apparent substitution rate (i.e., branch length) with fragmentation.

## Results

### Mitochondrial genome fragmentation is widespread among Amblycera

We assembled mitogenomes from 90 samples across 53 genera (89 species) of Amblycera (S1 Table), finding evidence for circularity in most chromosomes (186 out of 197, S3 Table). We were able to identify all protein-coding genes in the majority of samples (78, S2, S7-S21 Tables). Our phylogenomic analysis based on 2395 nuclear orthologs provided a well-resolved and supported tree for Amblycera, and there were only a few differences between the concatenated and coalescent trees, mostly involving rearrangements among some families (S1-S3 Figs). Although many species had single circular mitochondrial chromosomes, there were also many instances of mitogenome fragmentation (S4, S5 Figs), and some of these fragmentation events occurred over short timescales (< 5 Mya; S4, S5 Figs).

We found significant phylogenetic signal for mitogenome fragmentation within Amblycera (D-value = 0.281, P = 0.001, S13 Fig), consistent with a Brownian motion model (P = 0.18). Among the six currently recognised families of Amblycera (Boopiidae, Gyropidae, Laemobothriidae, Menoponidae, Ricinidae, Trimenoponidae; [26]), all except Gyropidae contain samples with single-chromosome mitogenomes (Fig 1). In particular, all samples from Ricinidae (6 spp.) and Boopiidae (2 spp.) possess single-chromosome mitochondrial genomes with largely consistent gene orders. Notably, most members of Ricinidae, the sister group to all other Amblycera, retain the gene order *atp8-atp6-cox3*, also seen in free-living book lice [9]. The largest family, Menoponidae, which exclusively inhabits birds, displays a high degree of mitogenome structural variation, ranging from single chromosome to highly fragmented across multiple clades (Fig 1). The families Gyropidae and Trimenoponidae, both exclusive to mammals, form a clade in our analysis, yet they are not mutually monophyletic. Within these families, mitogenome structure varies from single-chromosome (in *Trimenopon* and *Chinchillophaga*) to highly fragmented with up to nine mitochondrial chromosomes (in *Macrogyropus*).

**Fig 1.**
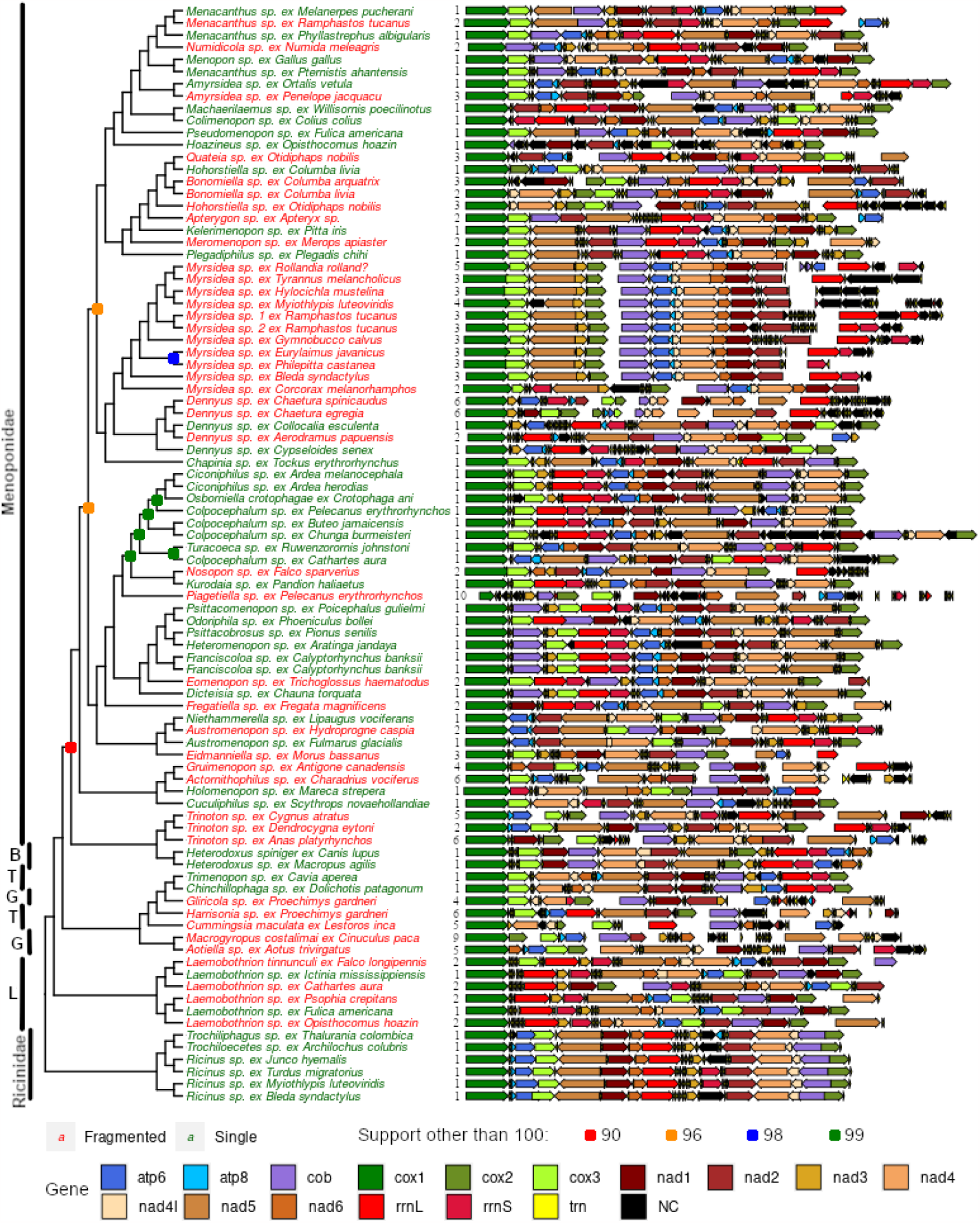
Phylogeny and pattern of mitogenome organisation across Amblycera. Chromosomes have been arbitrarily linearised, starting with *cox1* for consistent directionality and to facilitate comparison. Numbers next to the mitogenome plots indicate numbers of fragments. Gaps between genes indicate separate mitochondrial fragments. Taxa with fragmented mitogenomes are highlighted in red, and those with single-chromosome mitogenomes in green. All branches supported by 100% ultrafast bootstrap, except those indicated with coloured symbols at node. Abbreviations of amblyceran families: B – Boopidae; T – Trimenoponidae; G – Gyropidae; L – Laemobothriidae.

Our samples included multiple species within several amblyceran genera, allowing us to investigate variation in mitogenome structure among closely related taxa. In particular, we observed transitions from single-chromosome to fragmented mitogenomes within the genera *Menacanthus, Amyrsidea, Dennyus, Austromenopon*, and *Laemobothrion* (Fig 1). Moreover, we observed changes in the number of fragments between species within *Myrsidea* and *Trinoton* (Fig 1). Consistent with previous studies (e.g., [7, 19]), we found that gene order (Fig 1, S2 Table), even for single-chromosome genomes, was massively rearranged between lineages. However, in a few instances, the gene order remained stable among single-chromosome taxa, particularly within Ricinidae and Laemobothriidae (Fig 1, S2 Table). Our data suggest a possible trend towards increasing fragmentation within the genus *Myrsidea*. The sister taxon (*Myrsidea* sp. ex *Corcorax melanorhamphos*) to the rest of *Myrsidea* possesses two chromosomes, while the remaining species of *Myrsidea* have three chromosomes, or with some more derived species exhibiting even four or five chromosomes. This pattern suggests a gradual increase in the number of fragments over the course of the evolution of *Myrsidea* (Fig 1).

It is often assumed that once mitogenome fragmentation occurs, it is irreversible (e.g., [7]). Indeed, we found several highly supported cases where an ancestral state of a single chromosome transitions to a fragmented state (e.g., *Menacanthus, Nosopon, Piagetiella*, and *Eomenopon*). Despite this, our analysis did not reject the equal rates (ER) model for the evolution of mitochondrial fragmentation across Amblycera, and it was indeed the best-fitting model for this analysis (AIC = 109.8825, AICc = 109.928, AIC weight = 0.4197). When considering only Amblycera, this model suggests that the ancestral states of single versus fragmented have nearly equal likelihoods (S5 Fig). In this scenario, there are only three instances in the phylogenetic tree where a likely transition (>75% relative likelihood) from fragmented to single is inferred (*Hohorstiella* and two cases in *Laemobothrion*). However, for most taxa, the likelihood of the ancestral state does not strongly favour either state. One caveat is that our analysis does not take into account the genome organisation of more distant outgroups among free-living Psocodea, which is predominantly single, as is the case for most other insects. Therefore, it is likely that the ancestral state for Amblycera as a whole would be a single-chromosome in organisation (S4 Fig). The reconstruction that adopts this assumption (non-reversibility of fragmentation; AIC = 111.5266, AICc = 111.6646, AIC weight = 0.0040, S4 Fig) suggests at least 27 transitions from single-chromosome to fragmented mitochondrial genomes. While it remains to be seen if any definitive case of fragmentation reversal exists, further sampling within *Hohorstiella* and *Laemobothrion* could shed more light on this matter. Overall, fragmentation extensively varies across Amblycera, yet not to such an extent that it obscures overall evolutionary patterns.

### Rate of mitogenome fragmentation across Amblycera

The BAMM analyses provided evidence of significant changes in the rate of fragmentation over the amblyceran tree. The most probable solution in the analysis of the reconstructed rate of fragmentation (Fig 2) suggests seven rate shift events. In two cases (family Ricinidae and common ancestor of a group containing the *Colpocephalum* complex), the rate of fragmentation slowed down. The five cases in which the rate of fragmentation increased involved single genera (*Trinoton* and *Dennyus*) or terminal taxa (*Eomenopon, Piagetiella*, and *Nosopon*). The nine most frequent solutions of the analysis (S6 Fig) suggest between 5–8 rate shift events mainly positioned on the same branches as in the most probable solution.

**Fig 2.**
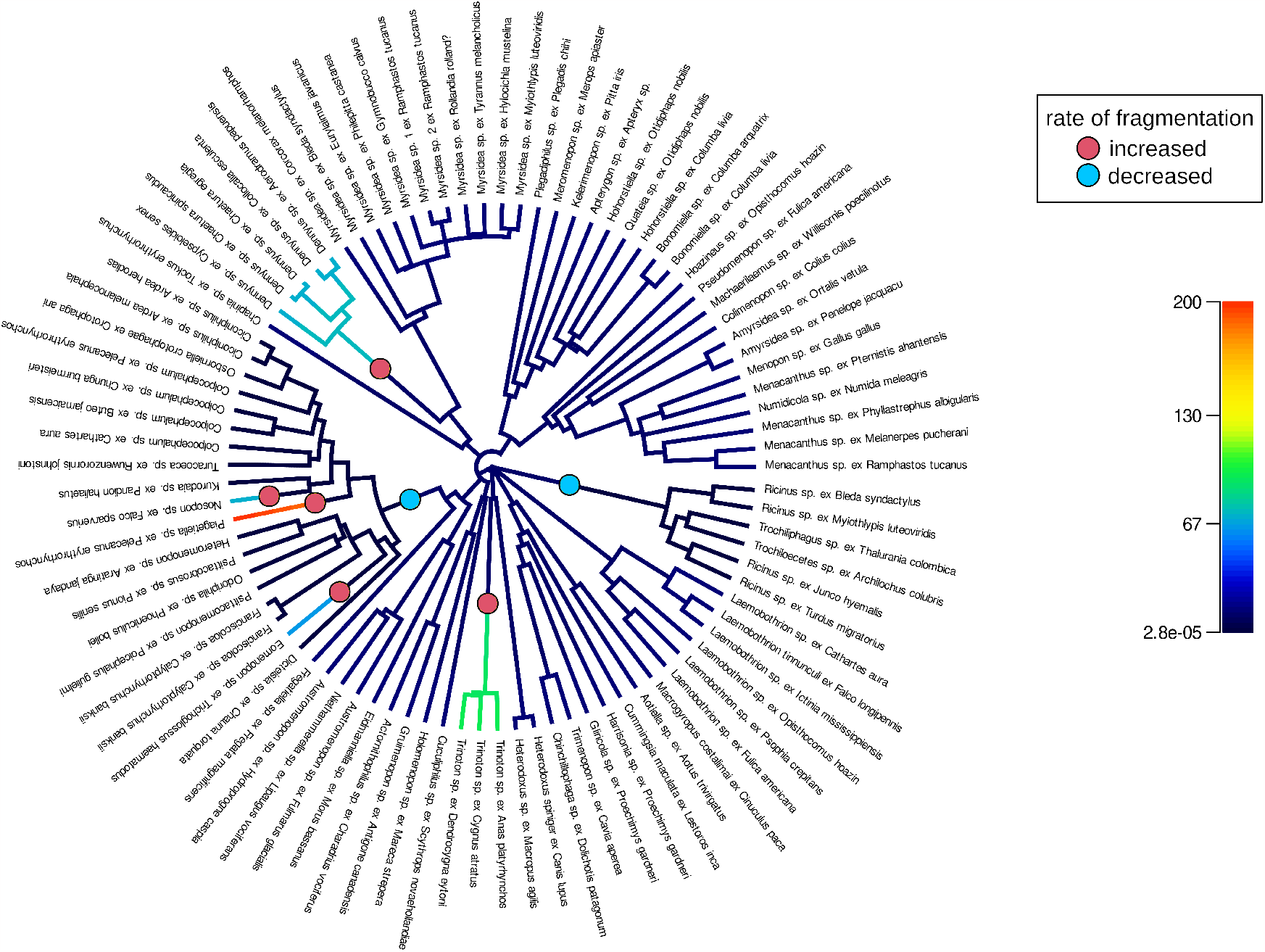
Changes in fragmentation rate in mitogenomes of Amblycera. Circles at the branches indicate rate shift events included in the optimal reconstruction (with maximum *a posteriori* probability). Rate increases with red, and rate decreases with blue. Overall reconstructed rate on each branch shown with color from low (dark blue) to high (red) as indicated by the scale bar.

### Comparison of substitution rates and gene rearrangements

We compared the lengths of both mitochondrial and nuclear branches in 19 Independent Matched Pair Comparisons within Amblycera (S7 Fig) between fragmented and non-fragmented lineages. For mitochondrial branch lengths, 14 of 19 pairs had a longer branch length in the fragmented lineage (Sign test *P* = 0.039; S7 Fig). This result indicates mitochondrial protein-coding genes evolve significantly faster in lice with fragmented mitogenomes compared to those with non-fragmented genomes. For nuclear branch lengths, 13 of 19 pairs had a longer branch length in the fragmented lineage (Sign test *P* = 0.108; S8 Fig), not quite significant.

Similarly, we compared gene rearrangements between fragmented and non-fragmented taxa for 19 Independent Matched Pair Comparisons within Amblycera (slightly different than the pairs for branch lengths comparison, S8 Fig). In 14 of 19 pairs, the fragmented mitochondrial genomes were more rearranged, compared to the reconstructed common ancestor, than non-fragmented genomes (Sign test *P* = 0.039). Although we did not manage to identify all tRNA genes in all the mitogenomes, the differences in gene rearrangements were so large (S6 Table) that even if we hypothetically placed them in the same positions as in their sisters, it would not affect the categorisation of which mitogenome is more rearranged. Consistent with Feng et al. [18] and Sweet et al. [7], the rRNA and tRNA genes were responsible for most gene rearrangements, having unstable positions between closely related species (e.g. genera *Amyrsidea, Bonomiella*; S2 Table).

### Fragmented mitogenomes of Amblycera are on average less AT biased

Comparison of the base composition (AT percentage) of single-chromosome mitochondrial genomes to fragmented genomes indicates that the AT composition of single-chromosome genomes is significantly higher overall (S9 Fig, S7 Table). Although there are some specific exceptions to these general patterns, the overall result is statistically significant. Although the AT content of fourfold degenerate sites is higher than that of other positions (S10 Fig), the AT content at fourfold degenerate sites from single mitogenomes is notably higher than that of sequences from fragmented mitogenomes (S11 Fig). The decrease in AT bias in fragmented mitogenomes is also observed at all other positions (S11 Fig). These findings align with previous studies [7, 10], reinforcing the hypothesis of differing mutational or selective biases in fragmented versus single-chromosome mitochondrial genomes. The differences in base composition across different protein-coding genes (S12 Fig, S8–S21 Tables) can be primarily attributed to disparities between fourfold degenerate sites and other partitions. In general, fourfold degenerate sites more likely reflect mutational biases rather than direct selection. These differences are especially evident in *cox1-3* and *cob* genes, which also exhibit the most conservation in spatial arrangement (Fig 1). We did not observe a significant correlation between chromosome length and AT content in fragmented mitogenomes.

### Phylogenomics clarifies phylogenetic relationships within Amblycera, uncovering paraphyly of some families

Both concatenated and coalescent analyses strongly support the monophyly of Amblycera (100% support; S1, S2 Figs). Within Amblycera (Fig 1; S1, S2 Figs), our phylogenomic tree confirms the monophyly of the families Ricinidae and Laemobothriidae (Fig 1). Concerning the family Boopidae, we examined only two samples of one genus, so although they are sister taxa, fully testing the monophyly of this family would benefit from more taxon sampling. Both the concatenated and coalescent trees suggest that the families Gyropidae and Trimenoponidae are paraphyletic. These two families intertwine to form a single monophyletic clade of lice (Fig 1; S1, S2 Figs) parasitising Neotropical mammals, primarily rodents and marsupials [26]. The concatenated (S1 Fig) and coalescent (S2 Fig) trees differ in the positions of the genus *Trinoton* and the family Boopiidae. In the concatenated tree (S1 Fig), the genus *Trinoton* is sister to Boopiidae, rendering Menoponidae paraphyletic. However, in the coalescent tree (S2 Fig), Boopiidae is a sister to all other Amblycera, while *Trinoton* is a sister to the remainder of Menoponidae, making this latter family monophyletic. In both trees, the remainder of Menoponidae collectively forms a large monophyletic clade (Fig 1; S1, S2 Figs). At the generic level, our data also suggest some genera may not be monophyletic, including *Austromenopon, Menacanthus, Hohorstiella, Colpocephalum*, and *Ricinus* (Fig 1; S1, S2 Figs).

## Discussion

By leveraging an extensive dataset of 90 samples of lice (53 genera, 89 total species) in the parvorder Amblycera, and integrating mitogenome assembly with nuclear phylogenomics, we gained substantial insights into mitogenome fragmentation within a diverse group of parasitic lice. We observed that mitogenomes undergo fragmentation numerous times (potentially 27 or more), with some fragmentation events occurring as recently as a few million years ago (S3, S4, S5 Figs). In various genera represented by more than one sample, transitions from a single chromosome to a fragmented mitogenome (e.g. *Dennyus, Laemobothrion, Menacanthus*), as well as increasing fragmentation into a larger number of fragments (e.g. *Myrsidea, Trinoton*) were apparent (Fig 1). Despite the occurrence of fragmentation events over many branches in the tree, the fragmentation process shows some evidence of phylogenetic conservation. Certain louse groups appear to fragment more frequently and rapidly than others, and some broad clades are either entirely fragmented or entirely single-chromosome. Among the six recognised amblyceran families, fragmentation has evolved in at least four of these, although the monophyly of some families is not supported. The reconstruction of the state of the ancestral mitogenome from current data depends heavily on the model used (i.e. whether reversal of fragmentation is permitted in the model or not). However, given the general conservation of mitogenome organisation across insects and in the common ancestor of Amblycera and free-living bark lice, it is highly probable that the common ancestor of Amblycera had a single mitogenome chromosome, with several transitions to fragmented mitogenomes occurring over time. Although transitions from single chromosome to fragmented chromosomes seem the most likely, the hypothetical merging of chromosomes [27] through homologous and non-homologous mitochondrial recombination [28] has been suggested as a possible mechanism by which fragmented mitogenomes might merge back into a single chromosome. However, even under the ER model (S4 Fig), the majority of changes are reconstructed to be from single to fragmented chromosomes.

In terms of the size of mitogenome fragments, we observed two primary patterns: 1) a small fragment encompassing a few genes splits from an original singe-chromosome mitogenome (e.g., *Laemobothrion*; *Numidicola*; *Menacanthus*; *Meromenopon*, Fig 1 [10]); 2) a single-chromosome mitogenome disintegrates into multiple smaller fragments of more similar sizes (e.g., *Myrsidea, Dennyus*, Fig 1; a similar pattern observed in Ischnocera, [7]). These two patterns are not mutually exclusive, and they can co-occur within a single individual (e.g., *Quateia*, Fig 1). In addition, numerous mitogenomes possess a state intermediate to these extremes. Furthermore, the genes involved, fragment sizes, and number of fragments seem to display considerable variation among different fragmentation events. Some studies [12, 13] hypothesised that mitogenomes comprised of numerous small minicircles would evolve incrementally from a few larger fragments rather than directly from a single-chromosome genome. However, the mitogenomes of the genus *Dennyus* (Fig 1) and the mitogenome of *Piagetiella* (Fig 1) suggest that a “big bang” fragmentation event, when multiple small fragments appear at relatively fast rate (Figs 1, 2), might indeed occur in lice, leading to a transition from a single to several mitochondrial chromosomes. Additional sampling within these genera might, however, uncover extant lineages with intermediate fragment numbers.

This variation suggests that the rate of fragmentation might change across the tree. In some cases, mitogenome structure is stable for a long period of time, while in others, it appears to change rapidly. In Ricinidae, both genome structure and gene order remained stable for at least around 30 million years (Fig 1; S4, S5 Figs). In contrast, over this same timeframe within the family Laemobothriidae, there are multiple transitions in mitogenome structure (Fig 1; S4, S5 Figs). These observations are borne out by detection of significant shifts in the rate of fragmentation over the tree using BAMM MCMC analyses. Specifically, the rate of fragmentation in Ricinidae is inferred to have decreased compared to the rest of the tree (Fig 2). Increases in the rate of fragmentation are recovered for *Dennyus* and *Piagetiella* (mentioned above), as well as on three other branches (Fig 2). In the genus *Myrsidea*, a relatively consistent three-fragment structure remained unchanged for an extended period across much of the diversification of the genus (33-10 Mya). However, in more recently diverged lineages within *Myrsidea*, additional fragmentation took place (S4, S5 Figs), but all these changes occurred at a similar rate to the overall rate across the tree (Fig 2). Given the notable diversity of *Myrsidea*, containing nearly 400 described species [26], it is likely that novel mitogenome configurations may be discovered in other species within this genus. In the case of *Dennyus*, the closest relative to *Myrsidea* (Figs 1, 2; S1–S5 Figs), it seems that its ancestor may have possessed a single-chromosome mitogenome which underwent a recent fragmentation event (within 2–5 My), resulting in six small minicircles (Figs 1, 2; S4, S5 Figs), and this likely occurred at a faster rate than changes over the rest of the tree. However, intermediate states of fragmentation cannot be completely ruled out. Even though evidence suggests mitogenome structure typically remains stable within a given species (e.g. *Columbicola passerinae* [7], *Franciscoloa* sp. ex *Calyptorhynchus banksii* [this study]), we uncovered instances where considerable variation can occur within a single genus (e.g. *Laemobothrion, Menacanthus*), sometimes over less than five million years (e.g. *Dennyus*, S4, S5 Figs).

Variation in the rate of fragmentation is a prediction of the heteroplasmy model of mitogenome fragmentation. In this case, lineages with chromosomal heteroplasmy would be prone to gene inactivation on one of the chromosome copies, which would then require the retention of the other copy of the chromosome [7]. This cycle would facilitate further gene deletion (i.e. fragmentation) within each chromosome. Thus, we would expect lineages with a heteroplasmic chromosomal state to fragment faster than lineages without, which would be resistant to fragmentation. The recombination model does not necessarily predict variation in the rate of fragmentation, but rather that fragmentation should be related to recombination errors, possibly associated with the overall rate of mutation. However, if the overall rate of mutation varies across the tree, the recombination model might also predict variation in the rate of fragmentation. Indeed, we found a correlation between whether a genome is fragmented or not and overall substitution rate (as measured by branch length, and which is directly proportional to mutation rate, all else being equal), which might support this model. It is unclear in this case, however, whether variation in mutation rate might cause fragmentation, or whether fragmentation causes an increased mutation rate.

Specifically, changes in chromosome length are expected to result in changes in base composition. Since shorter fragments would spend less time in the single-stranded state, they would be less subject to deamination mutations (changes to A and T) during replication. Our results also reveal that fragmented mitochondrial genomes exhibit a lower percent AT content compared to non-fragmented (single) ones. Compared to Ischnocera [7], the average AT percentage in Amblycera is marginally higher, possibly due to a higher proportion of single-chromosome mitogenomes. Regarding the AT percentage in different gene partitions, the fourfold degenerate sites could offer valuable insights into base composition biases, as they are not subjected to selection on the amino acid composition. Despite being significant, the difference in AT percentage between fourfold and non-fourfold degenerate sites in lice (approximately 5–10%, S10, S11 Figs, [7]) is similar to the difference in other insects [29]. The disparities among different PCGs (S12 Fig) indicate that the most pronounced effects of base composition biases are evident in the *cox* genes (involved with Complex IV in oxidative phosphorylation), consistent with the relatively strong purifying selection in these genes [30]. Overall, it could be a change in equilibrium base composition that leads to longer apparent branch lengths in fragmented versus non-fragmented mitogenomes.

We also found significant evidence that taxa with fragmented mitogenomes have more gene rearrangements than their non-fragmented relatives in paired comparisons. This finding is consistent with both the heteroplasmy and recombination models of mitogenome fragmentation. However, it is also the case that even non-fragmented mitogenomes within Amblycera have many gene rearrangements compared to the ancestral insect [19]. It seems unlikely that these rearrangements could be caused by heteroplasmy alone. However, recombination with heteroplasmic chromosome copies could potentially facilitate these rearrangements, so it may be that both heteroplasmy and recombination are important together to fully understand mitogenome evolution in lice.

Concerning the initial drivers of fragmentation in lice, although relaxed selection is typically associated with parasitism and reduced morphology, it does not account for why other arthropod parasites (e.g., fleas, ticks) lack fragmented mitogenomes or why fragmented mitogenomes also appear in free-living animals. We did, however, find a trend that lineages with fragmented mitogenomes also tended to have higher rates of substitution for nuclear genes (13 of 19 cases), although this was not statistically significant (*P* = 0.108). If this trend holds with additional sampling in the future, it might suggest that overall relaxed selection for mutation repair might be driving mitogenome fragmentation. Indeed, one hypothesis [12, 31] for the extremely fragmented mitogenomes of Anoplura is the absence of *mtSSB*, a nuclear gene that produces a protein (mitochondrial single-stranded DNA binding protein) targeted to the mitochondrion that stabilises mitochondrial DNA during the single-stranded replication stage. Lack of *mtSSB* could also be an explanation for the higher rate of DNA substitution in lineages with fragmented mitogenomes, because the mtSSB protein protects single-stranded DNA from mutations during replication. The presence of *mtSSB* in lice other than *Pediculus humanus* (Anoplura) is currently unknown; therefore, this hypothesis presents an avenue for further research. Overall, our dense taxon sampling within a single clade of lice provided an unprecedented number of phylogenetically independent comparisons of fragmented and non-fragmented mitogenomes to test important patterns of molecular evolution under this unusual phenomenon.

## Material & Methods

### Sequence data

We analysed the mitogenomes of 90 single specimens of amblyceran lice, a sample including all families, 53 genera, 89 species, the majority of host groups, and biogeographic regions across which Amblycera occur. Of these, 84 were newly sequenced for this study. We photographed individual specimens as vouchers and extracted total genomic DNA using a Qiagen QIAamp Micro Kit, with a 48-hour initial incubation [32]. We then prepared libraries from these extractions with a Hyper library kit (Kapa Biosystems) and sequenced them on an Illumina NovaSeq 6000 with 150bp paired-end reads [32]. We identified the vouchers to the genus level based on morphology using illustrations and keys [26, 33]. We also included data from six additional amblyceran species analysed by Sweet et al. [10] from NCBI SRA [16]. We conducted a quality check on the raw data from all 90 samples using FastQC v0.11.9 (https://www.bioinformatics.babraham.ac.uk/projects/fastqc/) and trimmed the reads using BBDuK from the BBMap package (https://sourceforge.net/projects/bbmap/, setup ktrim=r k=23 mink=7 hdist=1 tpe tbo maq=10 qtrim=rl trimq=35 minlength=35). We trimmed adapters automatically and manually trimmed the 5’ and 3’ ends using the forcetrim= argument.

### Mitochondrial genome assembly and annotation

To avoid excessive coverage of the mitochondrial genome (which can lead to assembly errors) and decrease the computational demand, each sequence read library was subsampled for 2 million reads of Read1 and the corresponding Read2 (4 million total reads). The mitochondrial genomes were first assembled using MitoFinder v1.4.1 [34] with MetaSPAdes as the assembler. From the MetaSPAdes results, contigs similar to the reference (i.e., concatenated nucleotide sequences of previously published mitogenomes; [10, 14]) were selected using TCSF v2.7.1 [35] with default parameters. From the TCSF results, contigs with high coverage (typically exceeding 100X) and a cumulative length of at least 15 kb were manually selected. These contigs were tested for circularity in Simple-Circularise (https://github.com/Kzra/Simple-Circularise) and AWA [36], searching for *k*-mers up to 40 bp long, mapped on full trimmed reads without subsampling. Manual inspection of gene overlap and sequence similarity, along with the AWA results, further validated circularity (S3 Table).

If the assembly failed to provide circular contigs encompassing all mitochondrial genes, the assembly procedure was repeated with subsampling increased to 8 million, and then 20 million total reads if necessary, to obtain sufficient coverage. For subsamples that produced successfully circularized contigs, we also tested increasing the number of reads for these libraries. In all of these cases (52), the assemblies with the higher subsample of reads were identical to those obtained with the lower subsample, suggesting that incrementally increasing the subsample of reads is an appropriate strategy both to avoid excessive coverage, but also to obtain sufficient coverage to produce accurate assemblies. Using MITOS2 [37], we annotated the contigs and identified genic regions overlapping the 3’ and 5’ ends. To verify annotation of protein coding genes (PCGs) and find PCGs not located by MITOS2, we manually searched open reading frames (ORFs) identified by ORF Finder, part of Sequence Manipulation Suite, [38], and visually inspected ORFs in Jalview v2.11.2.0 [39].

### Phylogenomics, dating, and ancestral state reconstruction

For phylogenomic analysis of Amblycera, we used whole genome sequencing data of 90 amblyceran samples and 29 outgroup taxa from Ischnocera, Trichodectera, Rhynchophthirina, Anoplura, and free-living Psocodea [40]. For this analysis, reads were trimmed for adaptors and quality (phred score < 30) using *fastp* v0.20.1 [41] and converted to aTRAM 2.1 [42] databases. We assembled a target set of 2395 single-copy ortholog PCGs from the human head louse (*Pediculus humanus*) for each genomic library using aTRAM, using *tblastn* and the AbySS assembler (iterations=3 and max-target-seqs=3000). The resulting sequences were annotated using Exonerate with the reference protein-coding sequences, exons aligned, and genes trimmed using established tools and parameters [32, 43]. We conducted a phylogenomic analysis on the concatenated alignment under maximum likelihood and using the GTR+G model in IQ-TREE 2 v2.1.2 [44]. Support was estimated using ultrafast bootstrapping (UFBoot2; [45] with 1000 replicates. We also performed a coalescent analysis in ASTRAL-III v5.7.4 [46] to account for gene-tree/species-tree discordance. Separate gene trees were inferred using IQ-TREE, based on the optimal models. Molecular dating analysis using the concatenated data set was conducted using Least Squares Dating (LSD2) in IQ-TREE 2 with calibration from previously published fossil and codivergence data (split between human and chimpanzee lice 5–7 Mya, split between the lice from Old World primates and Great Apes 20–25 Mya, the minimum age for Menoponidae of 44 Mya based on a fossil; [32, 43]) and root age 127.1 Mya [40]. Ancestral states of the mitogenome (single-chromosome or fragmented) were reconstructed over the dated tree using the *ace* function of the APE v5.4 R package [47] under various models: Equal-Rates (ER), All-Rates-Different (ARD), and a model that does not allow transition from fragmented to single mitogenome organisation (USR). The best model was selected using the corrected Akaike Information Criterion (AICc) and Akaike Information Criterion weight. The model fit was assessed with the *fitDiscrete* function in the GEIGER v2.0.7 R package [48], and the best model (ER; AIC = 109.8825, AICc = 109.928, AIC weight = 0.4197, S22 Table) revealed an almost 50% relative likelihood of fragmented ancestral amblyceran mitogenome. Given that more distant outgroups among Psocodea have non-fragmented mitogenomes, a fragmented ancestral state seems unlikely. Therefore, we also performed stochastic mapping with 1000 simulations for the USR models using the PHYTOOLS v0.7 R package [49].

To measure the strength of the phylogenetic signal of fragmentation, we calculated the D-statistic using the *comparative*.*data* and *phylo*.*d* functions of the CAPER v1.0.1 R package [50] over the dated amblyceran tree.

### Rate of fragmentation

To measure the rate of fragmentation and whether this rate changed over the tree, we analysed the dated concatenated tree in BAMM using Markov chain Monte Carlo (MCMC) simulations [51]. Although this program has become a subject of critique [52], this controversy applies only to quantifying speciation and diversification, not to analyses of phenotypic trait evolution. To set priors for BAMM MCMC cycles and to visualise BAMM outputs, we used the BAMMtools R package [53]. We ran the entire BAMM analysis in three iterations, with the following priors as calculated by BAMMtools: expectedNumberOfShifts = 1.0, betaInitPrior = 0.0136706518546954, betaShiftPrior = 2.90363806597022, useObservedMinMaxAsTraitPriors = 1. The MCMC simulations were run for 10 million generations each, with sampling every one thousand generations. We ran four Markov chains, proposed a chain swap every 1,000 generations, and reset the acceptance rate calculation every 1,000 generations.

### Comparison of branch lengths

The many independent transitions between single versus fragmented mitogenome structure in Amblycera (see Results) provided an excellent opportunity to test for the correlation of other factors with mitogenome fragmentation. One main expectation is that fragmented lineages evolve more rapidly [18], which should lead to longer reconstructed branch lengths in fragmented versus non-fragmented taxa. We compared branch lengths, both nuclear and mitochondrial, of taxa with single-chromosome versus fragmented mitogenomes to test whether fragmented mitogenomes evolve faster than the single-chromosome ones. To obtain the nuclear branch lengths, we used the concatenated tree from the phylogenomic analysis. To obtain mitochondrial branch lengths, the amino-acid sequences of mitochondrial protein-coding genes were aligned using UPP [54] with the -M -1 argument to filter out fragmentary sequences. Individual gene alignments were trimmed in trimAl [55] with a gap threshold of 0.4, back-translated using PAL2NAL [56], and concatenated using AMAS [57]. With the concatenated mitochondrial alignment, we estimated mitochondrial branch lengths by constraining the well supported tree from the nuclear gene set (see Results) and using maximum likelihood based on the optimal model (-m MFP) in IQ-TREE 2 v2.1.2 [44].

To compare branch lengths, we used the Independent Matched Pair Comparison technique [58, 59]. Specifically, we used a combination of sister pair comparisons and matched non-overlapping independent paired comparisons to construct contrasts between fragmented and non-fragmented lineages. In cases where a taxon was sister to a clade containing more than one taxon with the relevant mitogenome architecture (as in analogous situations in [59]), we selected the taxon with the median branch length for that clade. In these cases we scored whether the branch length to the most recent common ancestor for each pair was longer for the fragmented or non-fragmented lineage and compared these fractions using a two-tailed sign test, which is conservative in that it considers only the direction of difference rather than the magnitude [60].

### Ancestral gene orders and gene rearrangements

Given the many independent comparisons available for fragmented and non-fragmented taxa, we were also able to test for a correlation between mitogenome structure and gene rearrangement. As with comparisons of branch lengths (above), we used Independent Matched Pair Comparisons [58, 59] to select independent comparisons between fragmented and non-fragmented lineages. These comparisons were constructed to compare whether the number of gene rearrangements, compared to a reconstructed common ancestor, was higher in fragmented versus non-fragmented lineages. In cases where a selection of more than one representative of a clade was possible, we selected the taxon with the number of annotated mitochondrial genes nearest to that of the other taxon to which the comparison was being made. Across the concatenated tree, we reconstructed ancestral mitochondrial gene orders using AGORA v3.1 [61]. In each pair, we compared the numbers of shared gene boundaries shared with the most recent common ancestor. The shared gene boundaries were identified manually, as described by Feng et al. [18]. Because the numbers of annotated mitochondrial genes often differed in each pair member, we divided the number of shared gene boundaries by the number of annotated genes. We then tested whether a higher fraction of lineages with fragmented mitogenomes also had more gene rearrangements than non-fragmented lineages using a two-tailed sign test [60].

### Nucleotide composition

We calculated the nucleotide composition of the mitogenome for six sub-datasets for each sample (all sites, coding regions, different codon positions for all three positions, and fourfold degenerate sites of concatenated PCGs) using *Bio* and *Bio*.*SeqUtils* packages of Biopython 1.80 [62]. We identified the fourfold degenerate sites using MEGA11 [63]. We performed statistical comparisons of AT content using the GGPUBR v0.40 R package [64], the *phylANOVA* function in the PHYTOOLS v0.7 R package [49], taking into account the concatenated amblyceran tree. Additionally, we calculated AT content for each PCG separately. We used the average AT percentage of different PCGs to test for differences between genes with the Wilcoxon rank sum tests in the rstatix v0.7.2 R package [65]. We also assessed the correlation between AT content and the length of mitochondrial chromosomes, fitting both linear and Phylogenetic Least Squares (PGLS) models to the total AT content and AT content of fourfold degenerate sites, taking into account the concatenated amblyceran tree. For fragmented mitogenomes, we tested both the average length of chromosomal fragments and their actual lengths. For PGLS, we employed the *pgls* function of the CAPER v1.0.1 R package with both Brownian and Pagel’s λ correlations.

## Acknowledgements

We thank B. Benz, T. Chesser, D. Clayton, K. Dittmar, R. Faucett, T. Galloway, A. Grossi, F. Madeira, J. Malenke, M. Meyer, R. Moyle, B. O’Shea, R. Palma, V. Piacentini, A. Saxena, M. Valim, T. Valqui, J. Weckstein, and R. Wilson for assistance in obtaining samples for this study. We thank S. Virrueta Herrera for assistance with DNA extraction. We thank the A. Hernandez and C. Wright at the University of Illinois Roy J. Carver Biotechnology Center for assistance with Illumina sequencing. We also thank CSIRC personnel (Universidad de Granada, Spain) for assistance and for providing computational resources (Albaicin supercomputer). We thank K. K. O. Walden for assistance with the submission of raw read files to NCBI. This work was also supported by the Ministry of Education, Youth and Sports of the Czech Republic through the e-INFRA CZ infrastructure (ID:90140).

## Data and code availability

Data associated with this study are available in the Supplementary material and NCBI SRA. Other data are available on reasonable request. The code used to analyse mitogenomes is available on GitHub (https://github.com/tomas-najer/Amblycera-mitogenomes). Additional data, including photos of vouchers and alignments of all individual genes, are available on Figshare (https://figshare.com/account/home#/projects/169538).

## Authors contribution

TN assembled and analysed the mitochondrial genomes, wrote the first draft, edited the manuscript, and obtained funding. JD performed the phylogenomic and dating analyses and edited the manuscript. AB and ADS provided project conceptualisation, supervision and edited the manuscript. OS provided a morphological determination of the lice and edited the manuscript. KPJ conceptualised and supervised the work, provided samples and sequences, edited the manuscript, and obtained funding.

