## Supplemental figures for "Mitochondrial genome fragmentation is correlated with increased rates of molecular evolution"

**S1 Fig. Concatenated maximum likelihood phylogenomic tree of Amblycera.** Based on a target set of 2395 protein-coding genes. Numbers associated with branches indicate ultrafast bootstrap support.

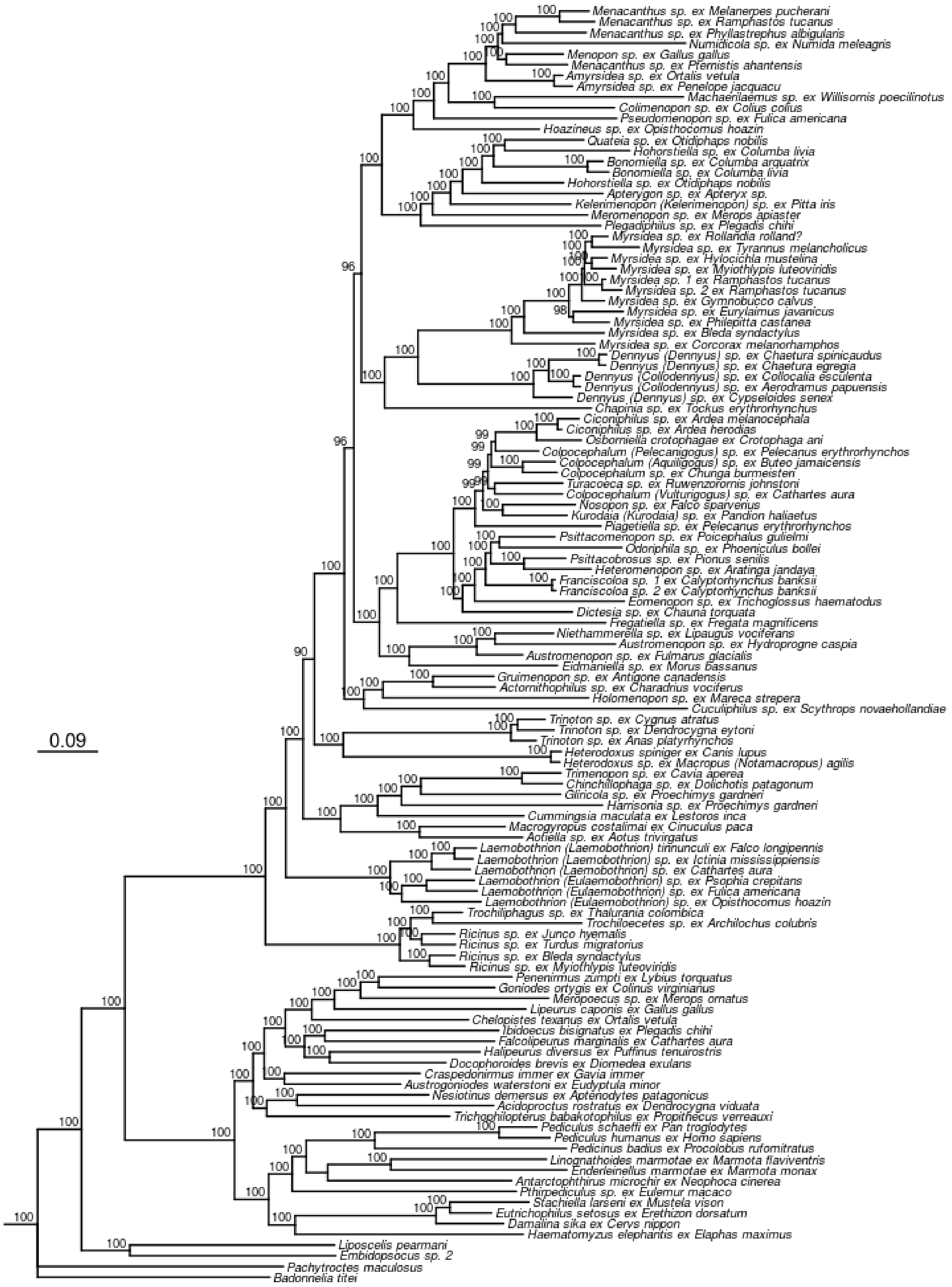

**S2 Fig. Coalescent tree of Amblycera.** Based on ASTRAL analysis of target set of 2395 protein-coding genes, combining individual gene trees into a species tree. Numbers associated with branches indicate local posterior probability.

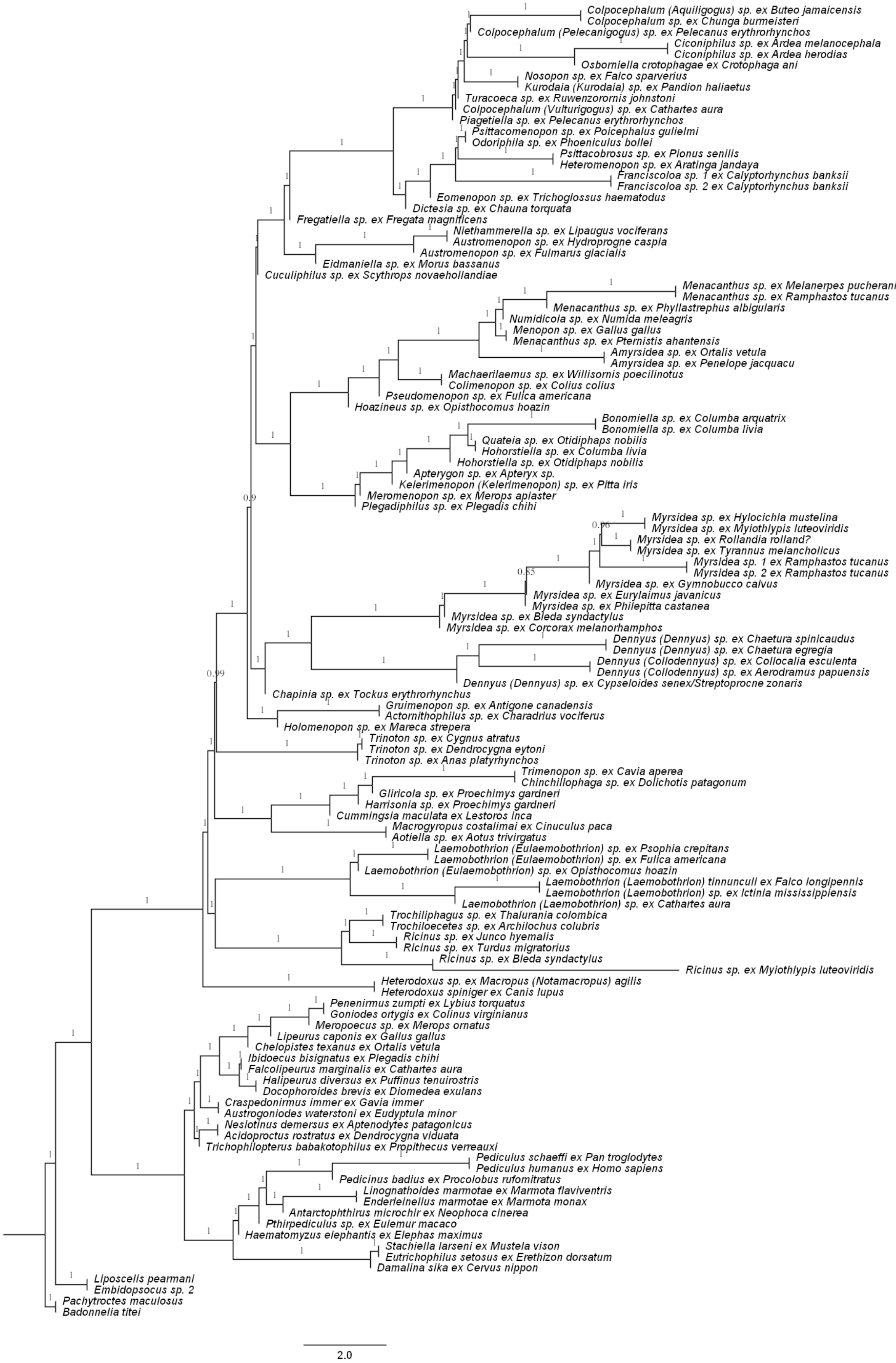

**S3 Fig. Dated phylogenomic tree of Amblycera.** Based on concatenated data set and a target set of 2395 protein-coding genes. Numbers at branches indicate 95% confidence intervals.

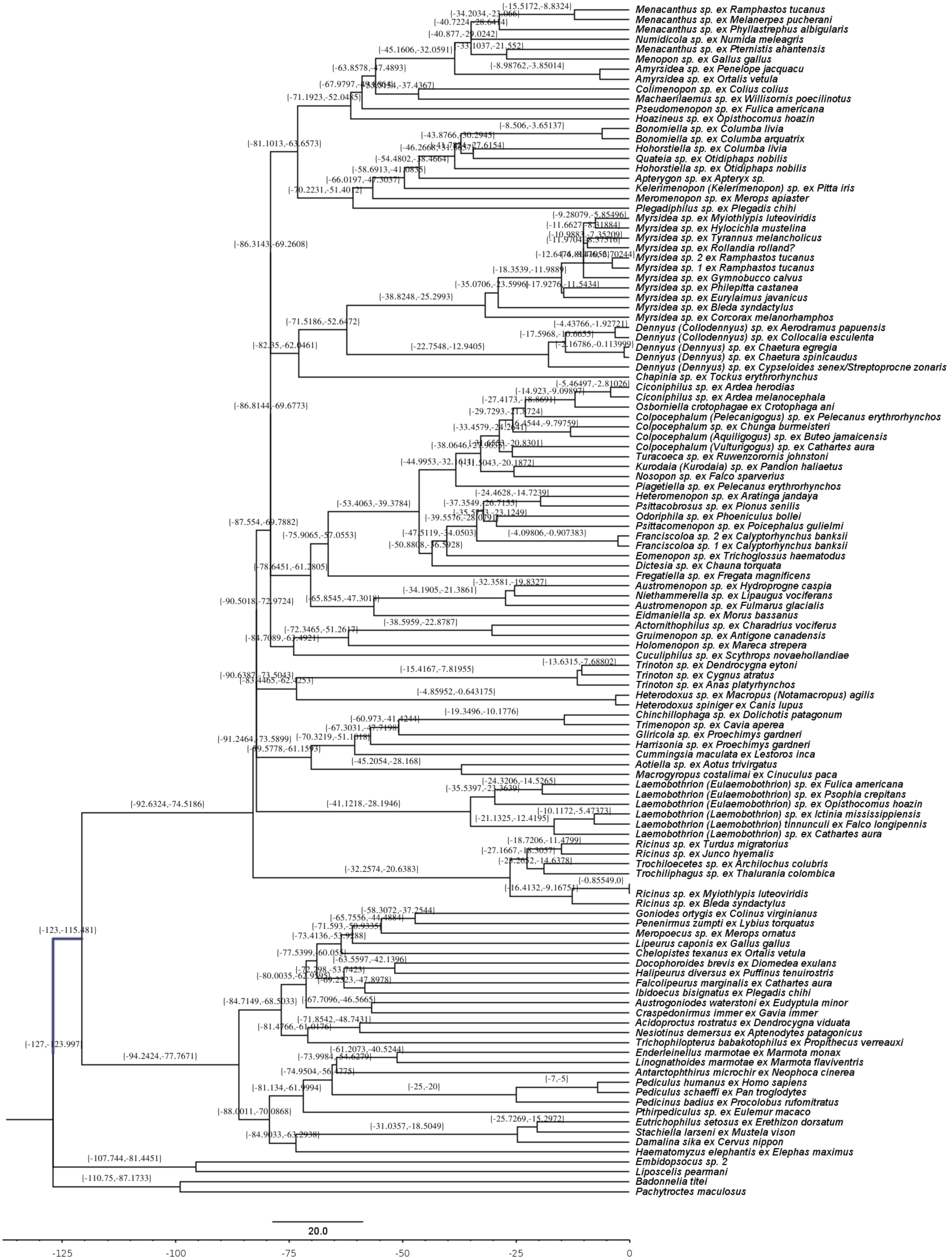

**S4 Fig. Dated phylogenomic tree with ancestral state reconstruction of mitogenome evolution in Amblycera under irreversible model.** Circles at the tips indicate mitogenome structure (single-chromosome vs fragmented). Pie charts at the nodes show the frequency distribution of reconstructed ancestral state after 1000 simulations of stochastic character mapping using an irreversible fragmentation model (USR). Time scale at bottom in million years ago (Mya).

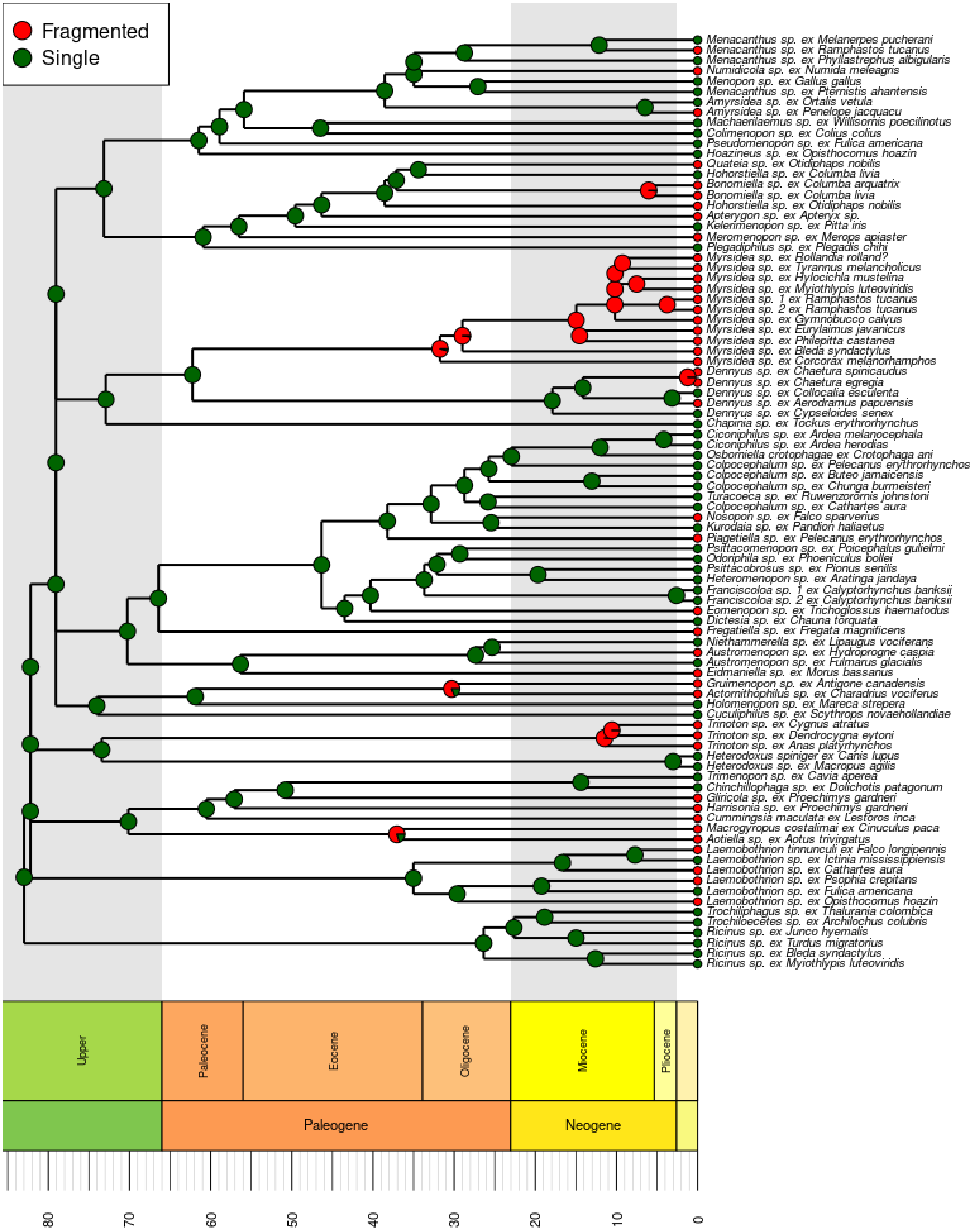

**S5 Fig. Dated phylogenomic tree with ancestral state reconstruction of mitogenome evolution in Amblycera under the Equal Rates (ER) model.** Pie charts at the nodes show the frequency distribution of reconstructed ancestral state after 1000 simulations of stochastic character mapping using with equal rates (ER) model. The ER model was best fitting according to the Akaike Information Criterion and corrected Akaike Information Criterion (AIC = 109.8825, AICc = 109.928). Circles at the tips indicate mitogenome structure (single-chromosome vs fragmented). Time scale at bottom in million years ago (Mya).

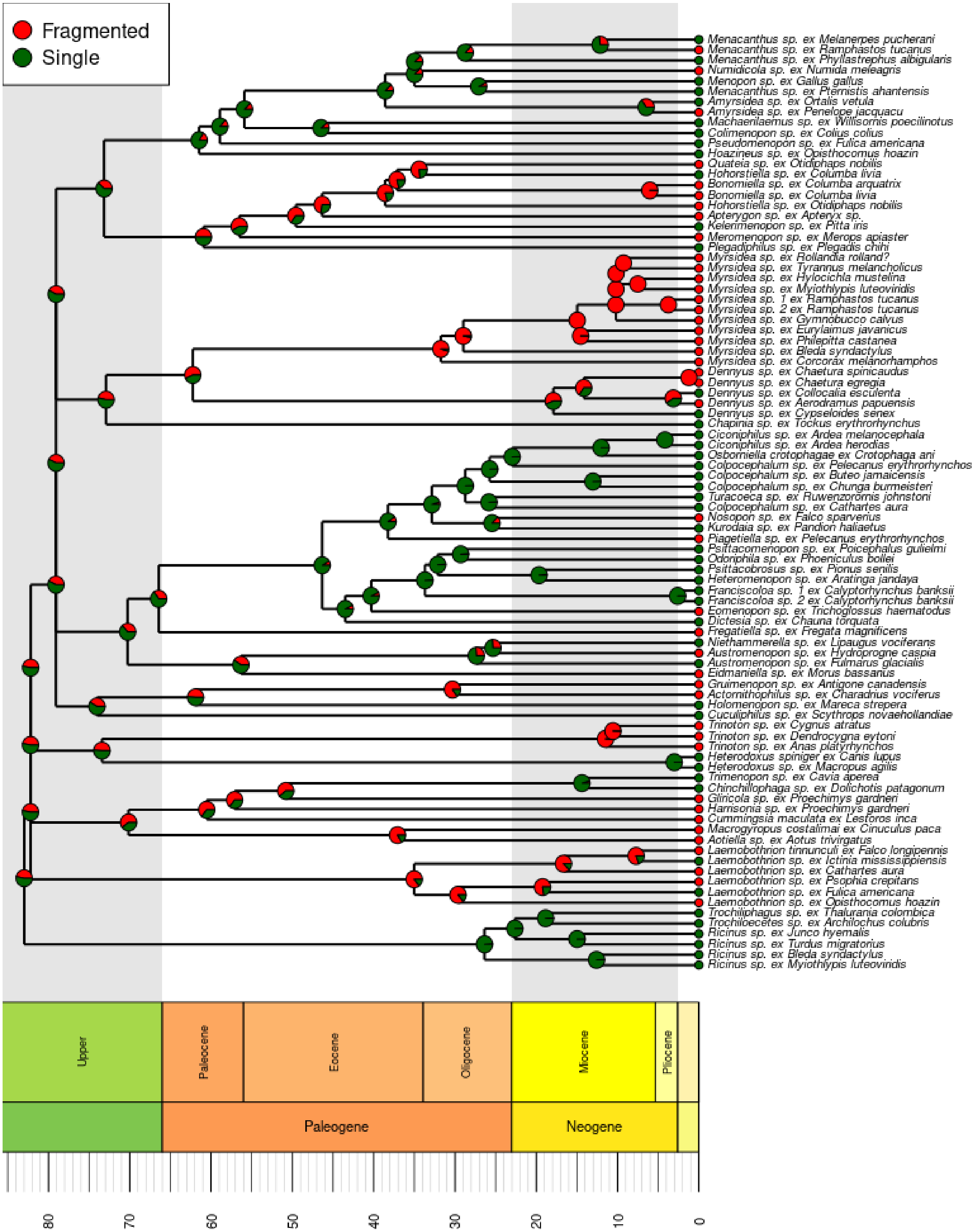

**S6 Fig. Bayesian credible set of configurations showing changes in fragmentation rate in mitogenomes of Amblycera.** Circles at the branches indicate rate shift events included in nine most probable reconstructions, numbers above the plots indicate posterior probabilities. Overall reconstructed rate on each branch shown with color, dark blue indicates the lowest, red indicates the highest.

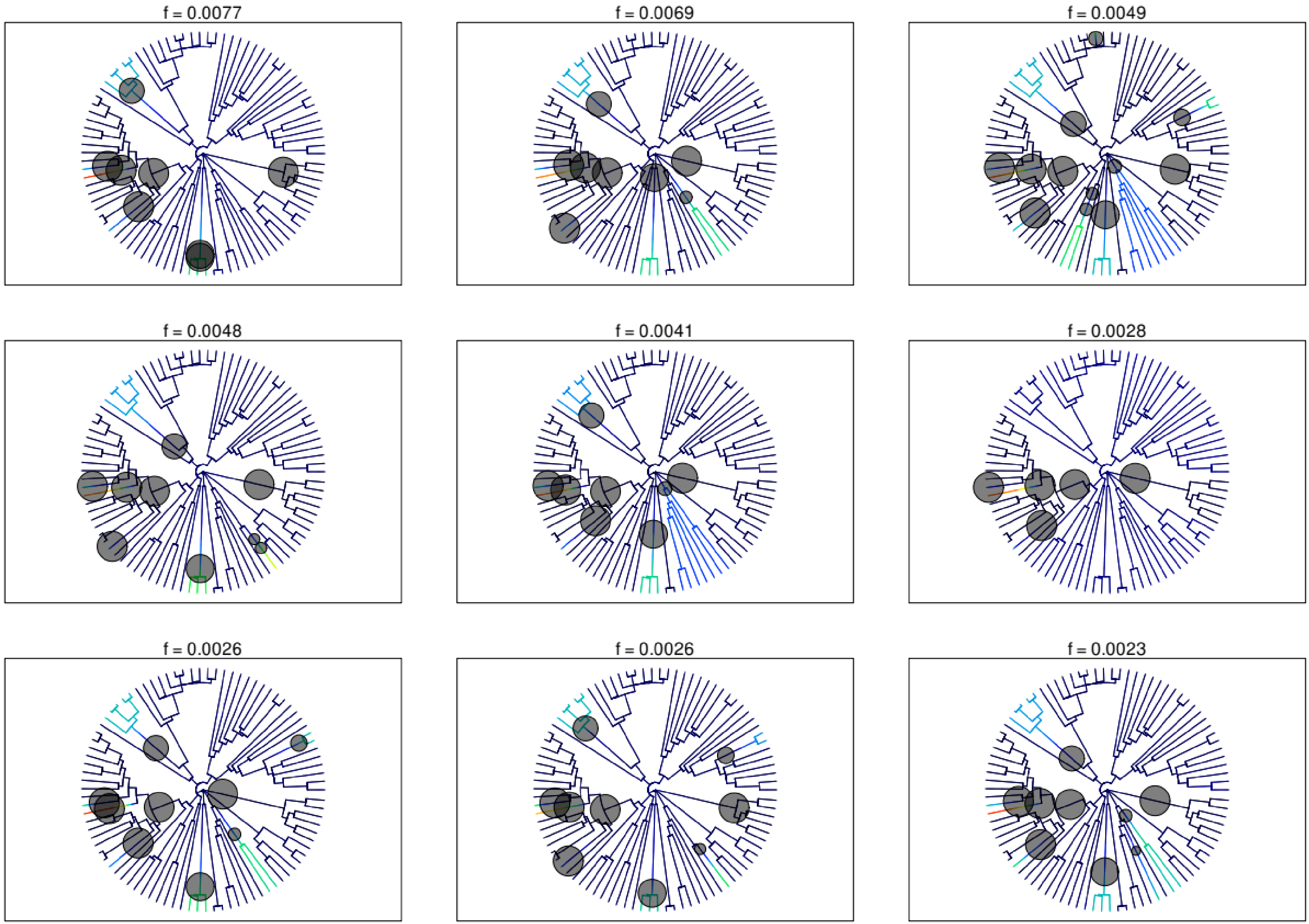

**S7 Fig. Pairs of Amblycera selected for comparison of branch lengths.** Circles at the tips indicate mitogenome structure (single-chromosome vs fragmented), and samples compared within a pair connected with orange line. Branch lengths computed from mitochondrial alignment, topology preserved from the nuclear concatenated tree.

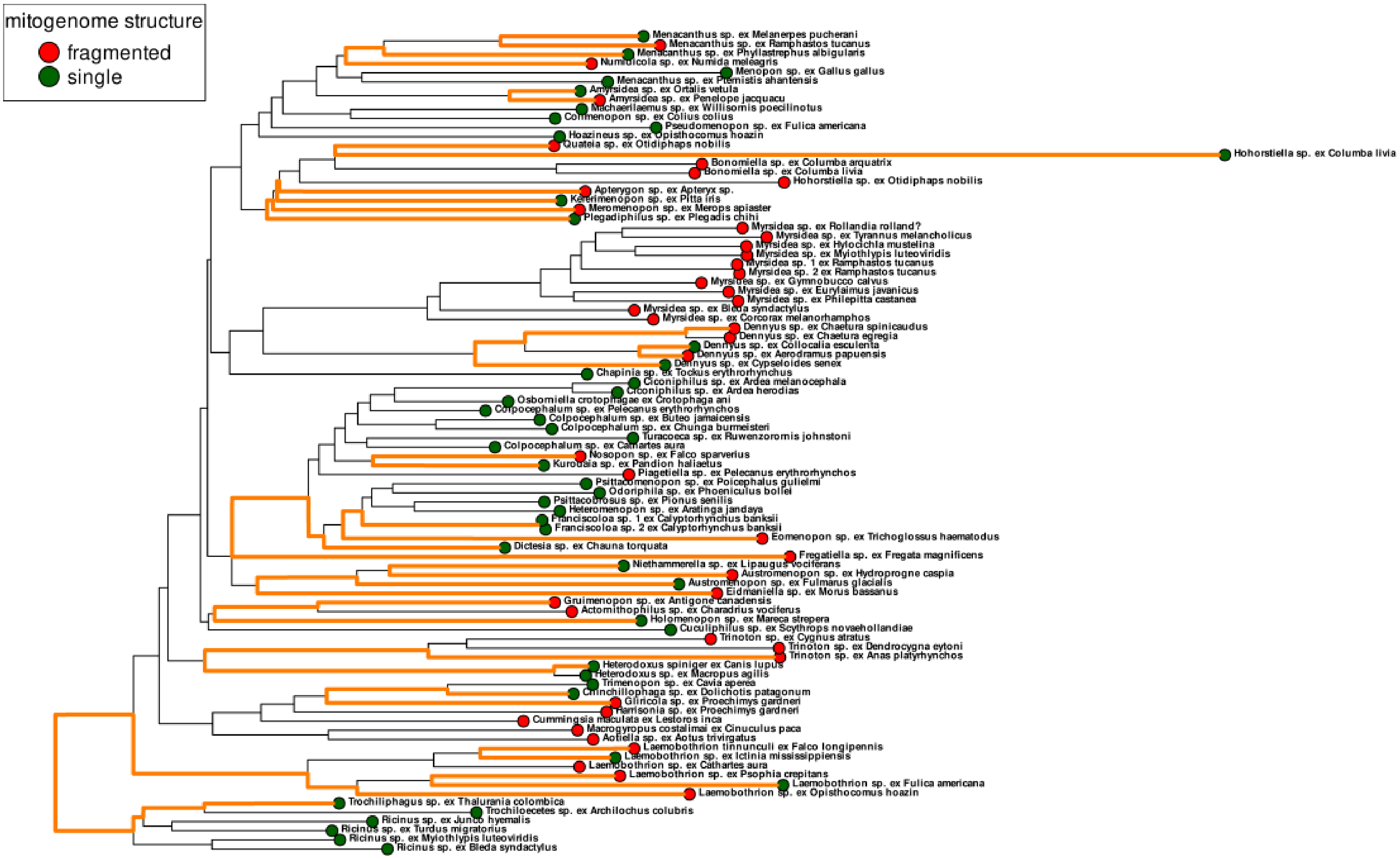

**S8 Fig. Pairs of Amblycera selected for comparison of gene rearrangements.** Circles at the tips indicate mitogenome structure (single-chromosome vs fragmented), and samples compared within a pair connected with blue line. Codes at the nodes indicate ancestral genomes, as in Tables S4 and S6. Both topology and branch lengths preserved from the nuclear concatenated tree.

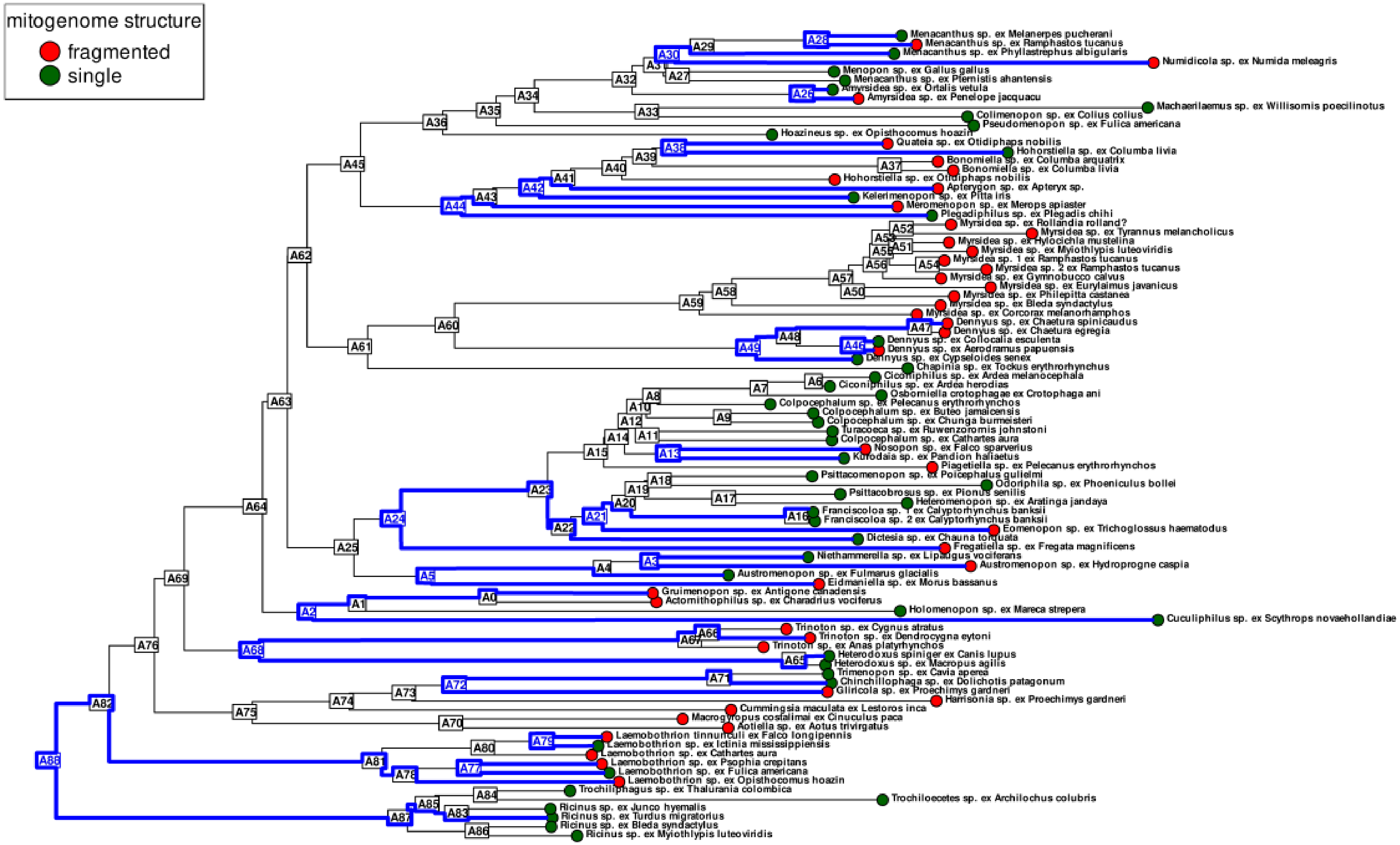

**S9 Fig. Comparison of AT content of fragmented (red) and single-chromosome (green) mitogenomes of Amblycera.** Values calculated from both coding and non-coding regions. Centerline – median; box limit – upper and lower quartiles; whiskers - interquartile range. The black dotted line shows the average percent AT content for insect mitogenomes available on NCBI GenBank (76.0%).

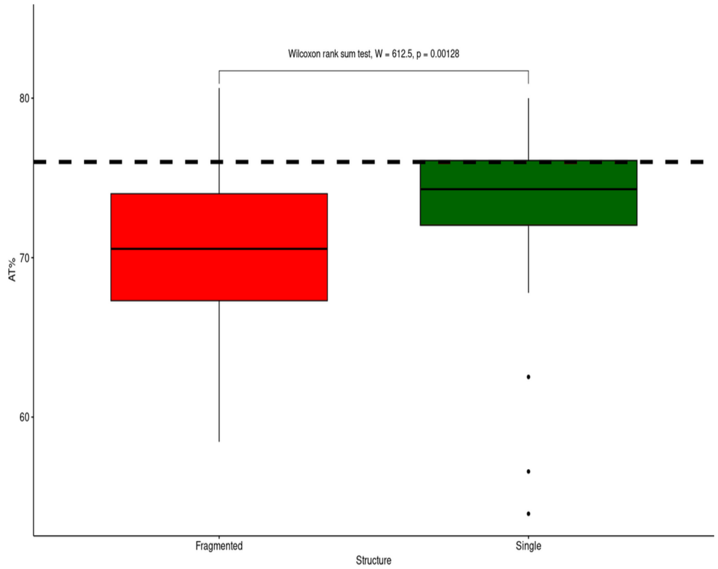


**S10 Fig. Percent AT content of different partitions of amblyceran mitogenomes.** Categories indicates values for entire sequences, coding regions, different codon positions, and fourfold degenerate sites. Centerline – median; box limit – upper and lower quartiles; whiskers - interquartile range. Significance based on Wilcoxon rank sum tests.

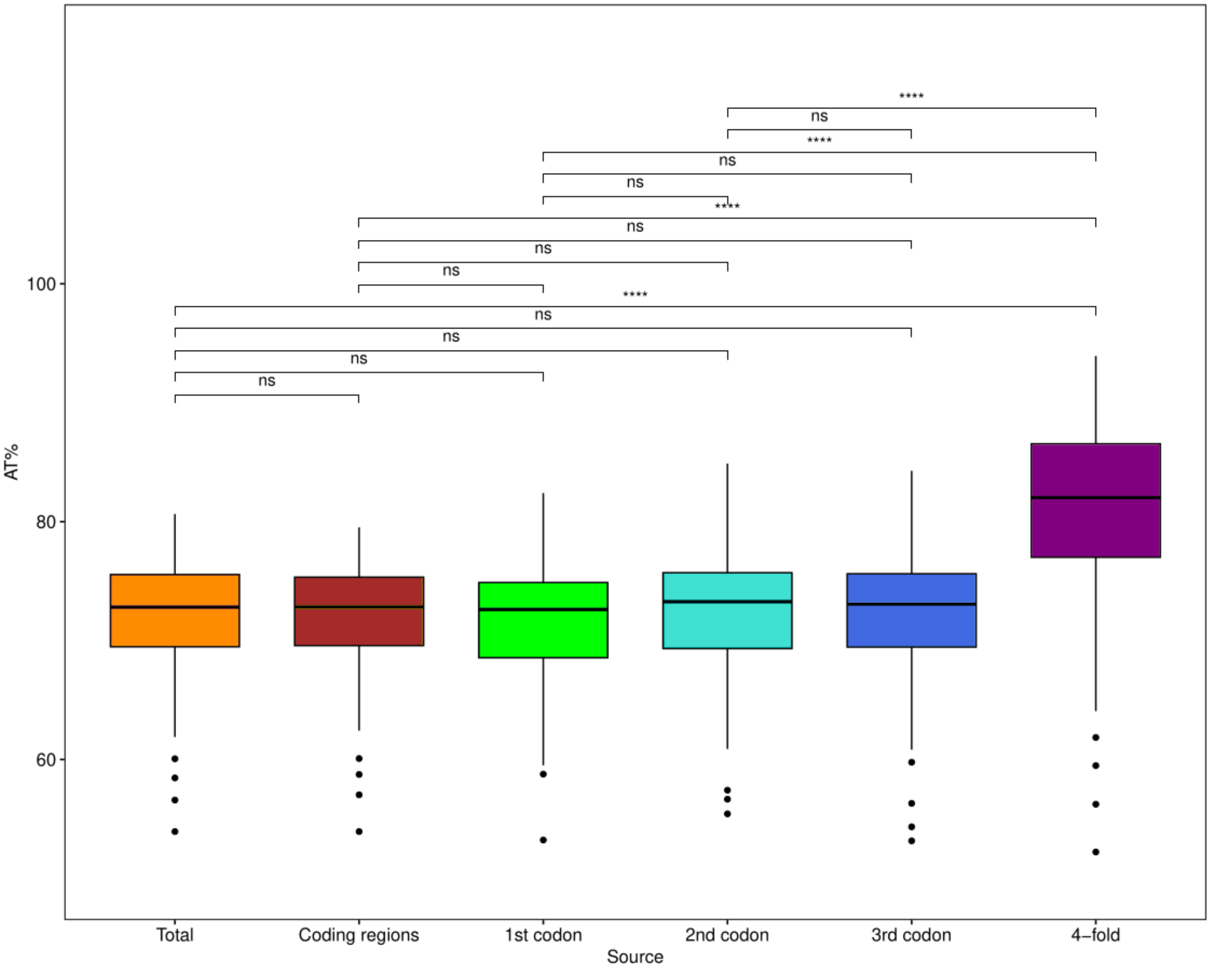

**S11 Fig. Percent AT content of different partitions of single-chromosome and fragmented amblyceran mitogenomes.** Categories indicate values for entire sequences, coding regions, different codon positions, and fourfold degenerate sites. Centerline – median; box limit – upper and lower quartiles; whiskers - interquartile range. The black dotted line shows the average percent AT content for insect mitogenomes available on NCBI GenBank (76.0%).

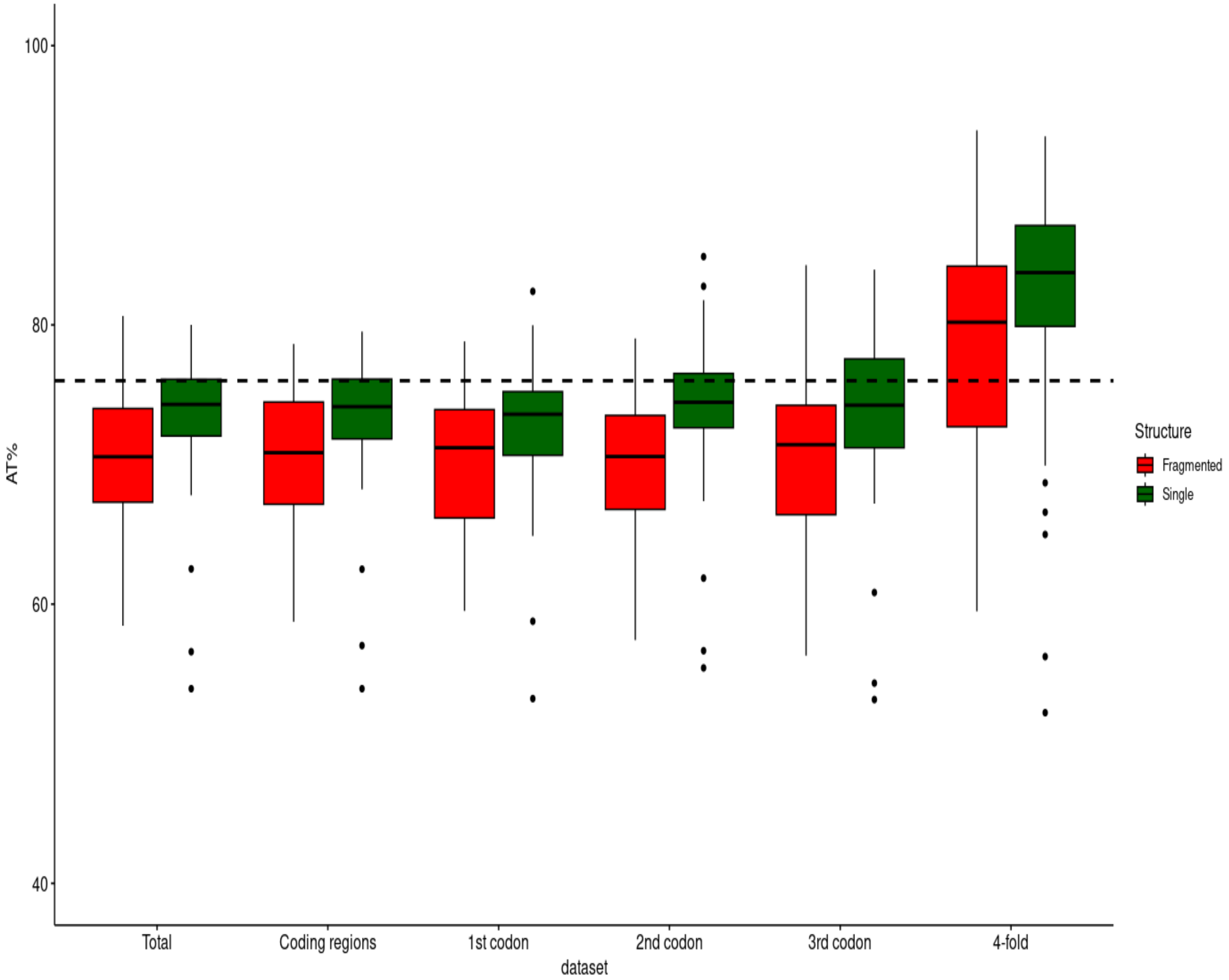

**S12 Fig. Percent AT content of different amblyceran protein-coding genes.** Showing differences between entire sequences and fourfold degenerate sites. Centerline – median; box limit – upper and lower quartiles; whiskers - interquartile range. The black dotted line indicates the average AT content for insect mitogenomes (entire sequences) available in NCBI GenBank (76.0%).

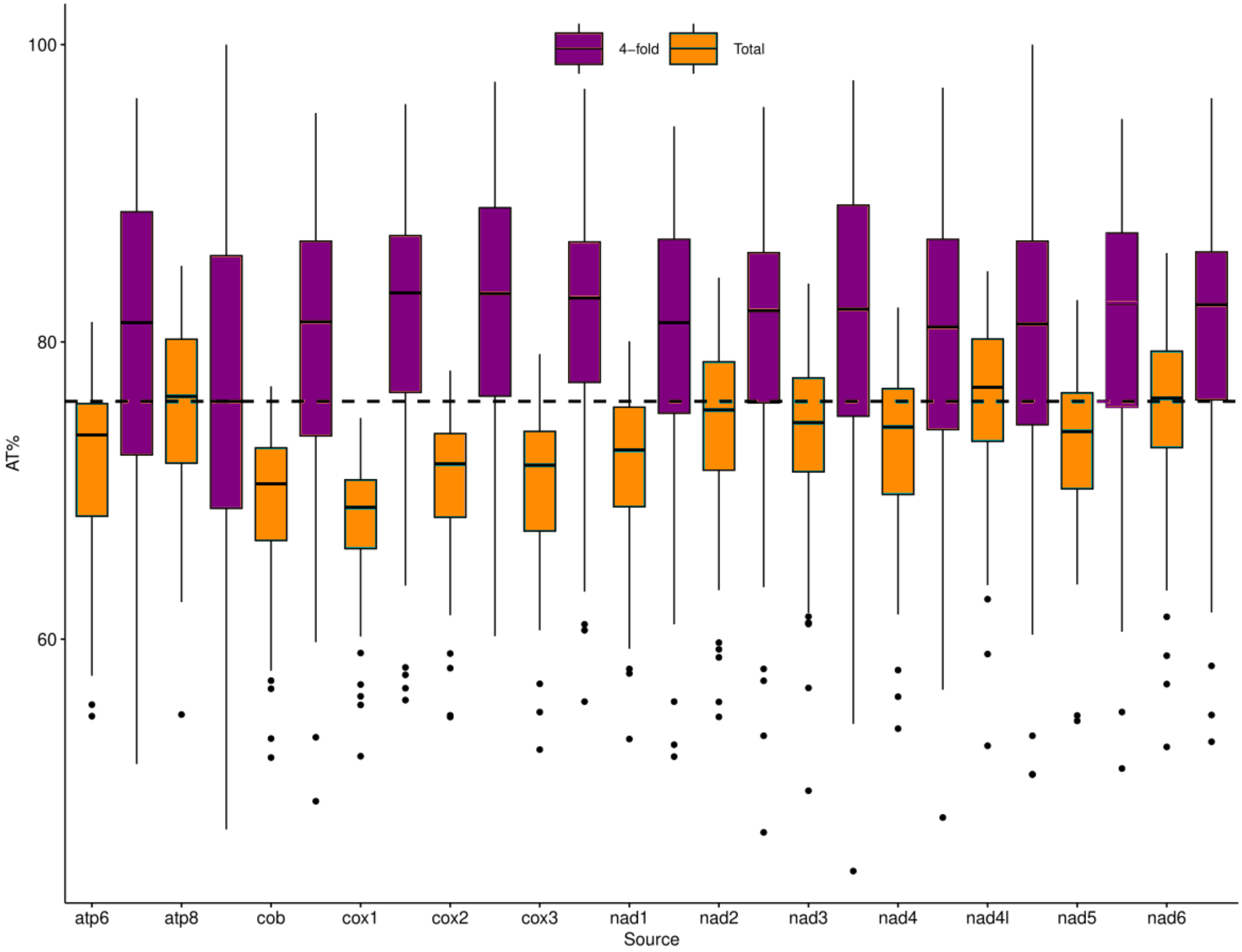

**S13 Fig. Phylogenetic signal of the mitogenome state of Amblycera (fragmented vs. single-chromosome).** Blue is values of D under simulation using Browning motion. Red is values of D under simulation with random phylogenetic structure. Black vertical line indicates value of D over actual tree (D = 0.281), *P* of E(D) under Brownian motion model = 0.177, *P* of E(D) under no phylogenetic structure model = 0.001.

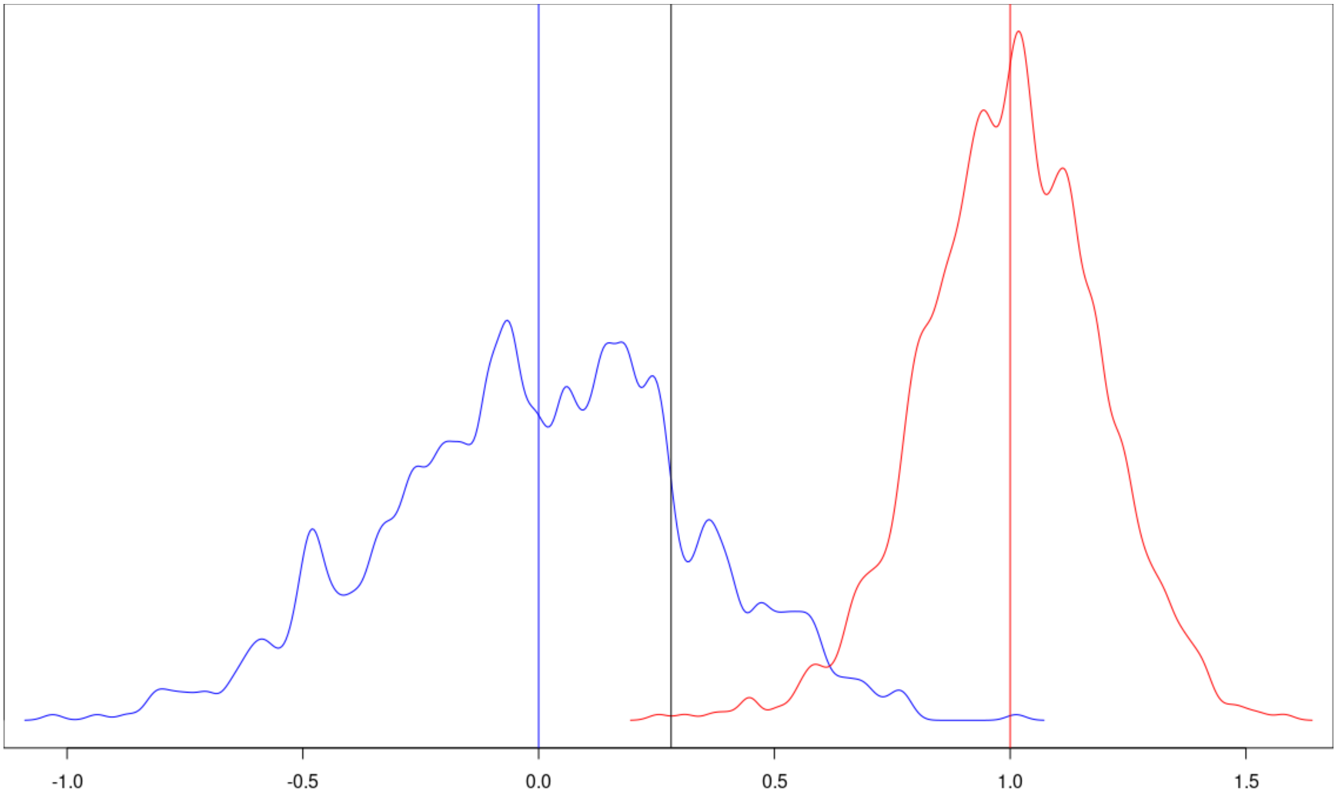


Estimated D : 0.2809441

Probability of E(D) resulting from no (random) phylogenetic structure : 0.001

Probability of E(D) resulting from Brownian phylogenetic structure : 0.177
